# An elusive endosymbiont: Does *Wolbachia* occur naturally in *Aedes aegypti*?

**DOI:** 10.1101/798736

**Authors:** Perran A. Ross, Ashley G. Callahan, Qiong Yang, Moshe Jasper, A. K. M. Arif, A. Noor Afizah, W. A. Nazni, Ary A. Hoffmann

**Affiliations:** Pest and Environmental Adaptation Research Group, Bio21 Institute and the School of BioSciences, The University of Melbourne, Parkville, Victoria, Australia; Institute for Medical Research, Kuala Lumpur, Malaysia

**Keywords:** *Wolbachia*, cytoplasmic incompatibility, *Aedes aegypti*, dengue

## Abstract

*Wolbachia* are maternally-inherited endosymbiotic bacteria found within many insect species. *Aedes* mosquitoes experimentally infected with *Wolbachia* are being released into the field for *Aedes-*borne disease control. These *Wolbachia* infections induce cytoplasmic incompatibility which is used to suppress populations through incompatible matings or replace populations through the reproductive advantage provided by this mechanism. However the presence of naturally-occurring *Wolbachia* in target populations could interfere with both population replacement and suppression programs depending on the compatibility patterns between strains. *Aedes aegypti* were thought to not harbor *Wolbachia* naturally but several recent studies have detected *Wolbachia* in natural populations of this mosquito. We therefore review the evidence for natural *Wolbachia* infections in *Ae. aegypti* to date and discuss limitations of these studies. We draw on research from other mosquito species to outline the potential implications of natural *Wolbachia* infections in *Ae. aegypti* for disease control. To validate previous reports, we obtained a laboratory population of *Ae. aegypti* from New Mexico, USA, that harbors a natural *Wolbachia* infection, and we conducted field surveys in Kuala Lumpur, Malaysia where a natural *Wolbachia* infection has also been reported. However, we were unable to detect *Wolbachia* infection in both the laboratory and field populations. Because the presence of naturally-occurring *Wolbachia* in *Ae. aegypti* could have profound implications for *Wolbachia*-based disease control programs, it is important to continue to accurately assess the *Wolbachia* status of target *Aedes* populations.

## *Wolbachia* infections in natural populations

*Wolbachia* are best known for their profound effects on host reproduction and more recently for their applied use in disease control programs. *Wolbachia* infect approximately half of all insect species but their prevalence varies widely between orders and genera (Weinert et al., 2015). Variation in infection also occurs within species, ranging from low frequencies to fixation (Charlesworth et al., 2019, Hilgenboecker et al., 2008). The prevalence of *Wolbachia* infections may be underestimated because infections can occur at low densities that are undetectable by conventional PCR (Mee et al., 2015). Multiple *Wolbachia* variants have been detected within the same species, such as in *Drosophila simulans* (Martinez et al., 2017) and *Culex pipiens* (Atyame et al., 2011). Superinfections, where multiple *Wolbachia* strains infect the same insect (Sinkins et al., 1995, Arthofer et al., 2009), also occur.

Although *Wolbachia* are maternally inherited, interspecific transfer may occur through parasitism (Heath et al., 1999, Ahmed et al., 2015), consumption of infected individuals (Le Clec’h et al., 2013, Brown and Lloyd, 2015), sharing a common environment (Huigens et al., 2004, Li et al., 2017) or other mechanisms. Successful horizontal transmission is likely to be rare, but *Wolbachia* can spread rapidly throughout populations once introduced (Kriesner et al., 2013, Turelli and Hoffmann, 1991). For *Wolbachia* to spread they must increase host fitness. *Wolbachia* infections may alter host reproduction to favor infected females over uninfected females, particularly through cytoplasmic incompatibility, which gives a frequency-dependent advantage to infected females (O’Neill et al., 1997). Cytoplasmic incompatibility results in fewer viable offspring in crosses between *Wolbachia*-infected males and uninfected females. *Wolbachia* may also provide fitness advantages through the protection of hosts against viruses (Teixeira et al., 2008, Hedges et al., 2008), nutritional provisioning (Brownlie et al., 2009), increased fertility (Dobson et al., 2002) or changes in life history (Cao et al., 2019).

Insects that are not naturally infected with *Wolbachia* may be amenable to infection experimentally. Novel *Wolbachia* infections have been generated through microinjection, where cytoplasm or purified *Wolbachia* from an infected donor is transferred to an uninfected embryo (Hughes and Rasgon, 2014). Deliberate transfers of *Wolbachia* between species are challenging and can take thousands of attempts to generate a stable line (McMeniman et al., 2009, Walker et al., 2011). But once an infection is introduced, *Wolbachia* infections have applications for pest and disease vector control since they can alter host reproduction and block virus replication and transmission (Hoffmann et al., 2015).

## Releases of novel *Wolbachia* infections for vector and disease control

There is increasing interest in deploying mosquitoes with experimentally-generated *Wolbachia* infections into the field for disease control. Over 25 novel *Wolbachia* infection types have been generated in mosquitoes through embryonic microinjection, mainly in the principal dengue vectors *Ae. aegypti* and *Ae. albopictus* (Ross et al., 2019b). Most of these infections induce cytoplasmic incompatibility and many also reduce the ability of their hosts to transmit viruses, making them desirable for field release. For mosquito species that are naturally *Wolbachia*-infected such as *Ae. albopictus*, novel infections can be generated either by first removing the natural infections with antibiotics (Suh et al., 2009, Calvitti et al., 2010) or by introducing the novel infection into an infected mosquito, resulting in a superinfection (Zhang et al., 2015, Suh et al., 2016). Different novel *Wolbachia* infections may be incompatible with each other (Ant et al., 2018) and the addition of *Wolbachia* strains to create superinfections can lead to unidirectional incompatibility, where females of the superinfected strain produce viable offspring following matings with males with any infection type, but superinfected males induce cytoplasmic incompatibility when mated with singly infected and uninfected females (Joubert et al., 2016).

Mosquitoes with novel *Wolbachia* infections are being released into the field for two main purposes: population replacement and population suppression. The objective of the former approach is to replace natural populations with mosquitoes possessing *Wolbachia* infections that interfere with virus transmission. This is achieved through the release of males that induce cytoplasmic incompatibility and females that transmit the *Wolbachia* infection and have reduced vector competence (Walker et al., 2011). Successful population replacement of *Ae. aegypti* with novel *Wolbachia* infections has been achieved in several countries (Hoffmann et al., 2011, Garcia et al., 2019, Nazni et al., 2019). Following releases in Australia and Malaysia, *Wolbachia* infections have maintained a stable, high frequency in most locations, coinciding with reduced local dengue transmission (O’Neill et al., 2018, Ryan et al., 2019, Nazni et al., 2019). Population suppression can be achieved through male-only releases of *Wolbachia*-infected males, resulting in cytoplasmic incompatibility with wild females. This was first demonstrated in 1967 in *Cx. pipiens* (Laven, 1967) by exploiting the natural variation in *Wolbachia* infection types between mosquitoes from different locations (Atyame et al., 2014). Other releases have used *Wolbachia* from a closely related species through introgression (O’Connor et al., 2012) and novel *Wolbachia* transinfections generated through microinjection (Mains et al., 2016, Zheng et al., 2019).

Both population replacement and suppression approaches rely on the novel *Wolbachia* infection types inducing cytoplasmic incompatibility with the resident mosquito population. Thus, the presence of natural *Wolbachia* infections in mosquitoes may interfere with disease control programs, making population replacement or suppression challenging or even impossible.

## Detections of *Wolbachia* in *Aedes aegypti*

*Aedes aegypti* is the principal vector of dengue virus and has been the focus of *Wolbachia*-based population replacement efforts, with releases of mosquitoes with novel *Wolbachia* infections now underway in over 10 countries (e.g. Nazni et al. (2019), Garcia et al. (2019), Hoffmann et al. (2011)). Until recently, *Ae. aegypti* was not thought to harbor *Wolbachia* naturally (Kittayapong et al., 2000), though it is clearly amenable to infection given the number of stable experimental infections generated in this species (Ross et al., 2019b). Evidence for horizontal gene transfer between *Wolbachia* and *Ae. aegypti* may reflect a historical infection (Klasson et al., 2009). The most comprehensive survey to date found no evidence for *Wolbachia* infection in *Ae. aegypti* through PCR assays on pools of mosquitoes, except in a single location where the experimentally-generated *w*Mel strain of *Wolbachia* had been released deliberately (Gloria-Soria et al., 2018). The lack of natural infection is advantageous for both population replacement and suppression programs because any cytoplasmic incompatibility-inducing *Wolbachia* infection should be unidirectionally incompatible with wild populations.

Coon et al. (2016) detected *Wolbachia* in *Ae. aegypti* collected from Florida, USA using 16S rRNA sequencing and multi-locus sequence typing. This discovery suggested that natural *Wolbachia* infections may occur in *Ae. aegypti*, with its occurrence perhaps being geographically restricted or at a low frequency in other populations. Since then, seven further studies have purported to detect *Wolbachia* in natural populations of *Ae. aegypti* (Table 1). These studies report variable infection frequencies in populations and identify infections from several *Wolbachia* supergroups. Most studies found that the infections detected were closely related to or identical to the *w*AlbB infection that occurs natively in *Aedes albopictus* (Coon et al., 2016, Balaji et al., 2019, Carvajal et al., 2019, Kulkarni et al., 2019), while other studies also detected *Wolbachia* from supergroups that do not normally occur within Diptera (Carvajal et al., 2019, Thongsripong et al., 2018). Most evidence is limited to molecular detection but some studies established laboratory colonies and have reported maternal transmission of *Wolbachia* (Kulkarni et al., 2019) and the loss of infection through antibiotic treatment (Balaji et al., 2019).

**Table 1.** Reports of natural *Wolbachia* infections in *Aedes aegypti*.

| Location | Collection date(s) | Evidence for infection | Infection frequency (n tested) | Supergroup | Reference |
| --- | --- | --- | --- | --- | --- |
| Jacksonville, Florida, USA | July 2014 | Molecular detection (16S rRNA sequencing, MLST) | Not specified | A and B | Coon et al. (2016) |
| Kuala Lumpur, Malaysia | Not specified | Molecular detection (*wsp*) | 25% (16) | Unknown | Teo et al. (2017) |
| Nakhon Nayok, Thailand | 2008 | Molecular detection (16S and 18S rRNA sequencing) | Not specified | C, others | Thongsripong et al. (2018) |
| Houston, Texas, USA | Not specified | Molecular detection (16S rRNA sequencing) | Not specified | Unknown | Hegde et al. (2018) |
| Tamil Nadu, India | August 2015 | Molecular detection (16S rRNA, *wsp*, *ftsZ*, MLST)<br>Electron microscopy qPCR across developmental stages and tissues<br>Removal through antibiotic treatment | Not specified | B | Balaji et al. (2019) |
| New Mexico and Florida, USA | 2016, 2017 | Molecular detection (PCR, LAMP)<br>Maternal transmission | 44.8% (194) | B | Kulkarni et al. (2019) |
| Manila, Philippines | May 2014-January 2015 | Molecular detection (*wsp*, 16S rDNA) | 11.9% (672) | A, B, C, D and J | Carvajal et al. (2019) |
| Panama | Not specified | Molecular detection (16S rRNA sequencing) | 0.2% (490) | Unknown | Bennett et al. (2019) |

Similar to *Ae. aegypti, Anopheles* mosquitoes (which transmit *Plasmodium* parasites that cause malaria) were also thought to be uninfected by *Wolbachia*, though several recent studies have detected *Wolbachia* in this genus (Baldini et al., 2014, Jeffries et al., 2018, Ayala et al., 2019). In a critical analysis of studies in *Anopheles gambiae*, Chrostek and Gerth (2019) assert that the evidence is currently insufficient to diagnose natural infections in this species. We highlight similar issues with detections of *Wolbachia* in *Ae. aegypti* but also discuss the potential implications for disease control if *Wolbachia* do occur naturally in this species.

## Potential implications of natural *Wolbachia* infections for releases of novel infections

The presence of natural *Wolbachia* infections may influence compatibility patterns between mosquitoes with the novel *Wolbachia* infection and the natural population. These patterns are summarized in Figure 1, although crossing patterns in nature are likely to be more complex. Natural *Wolbachia* infections can have heterogeneous densities and frequencies in populations (Calvitti et al., 2015), making compatibility patterns hard to predict. Crosses may differ in the strength of incompatibility in different directions (O’Neill and Paterson, 1992, Sinkins et al., 1995, Joubert et al., 2016) and there are also environment-dependent effects on cytoplasmic incompatibility including adult age (Kittayapong et al., 2002b) and temperature (Ross et al., 2019a). The presence of *Wolbachia* superinfections also increases the number of potential compatibility patterns (Dobson et al., 2004).

**Figure 1.**
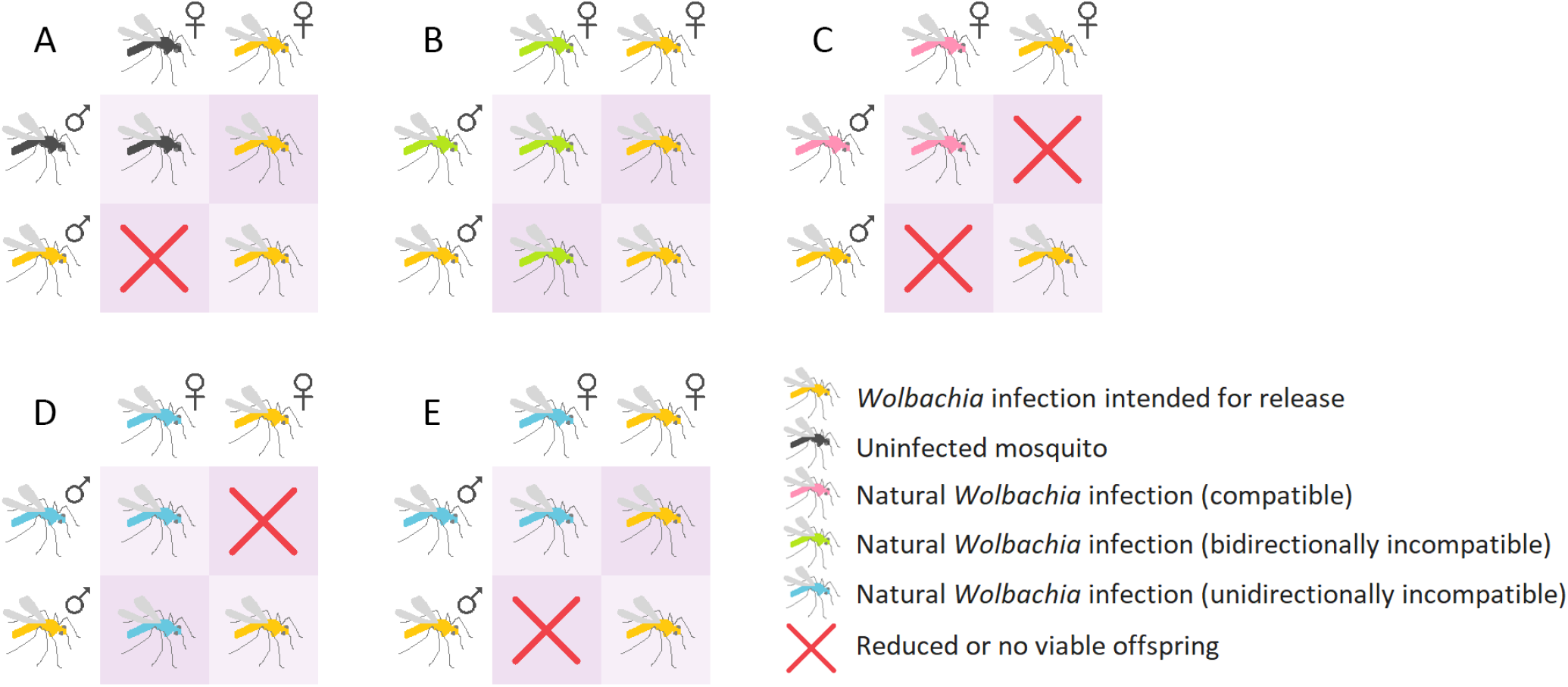
Potential crossing patterns between mosquitoes with novel *Wolbachia* infections that induce cytoplasmic incompatibility and mosquito populations with or without the presence of natural *Wolbachia* infections of different crossing types. (A) Crosses between mosquitoes with a novel *Wolbachia* infection and uninfected mosquitoes result in unidirectional cytoplasmic incompatibility. (B) When novel and natural *Wolbachia* infections exhibit the same crossing type, no cytoplasmic incompatibility occurs. (C) Bidirectional incompatibility occurs when novel and natural *Wolbachia* infections exhibit different crossing types. (D-E) Unidirectional cytoplasmic incompatibility may occur in favour of the natural (D) or (E) novel infection. These situations are most likely when the natural (D) or novel (E) infection is a superinfection, where one strain is compatible with the single infection but the other is not.

With most novel infections generated in *Ae. aegypti*, the release of *Wolbachia*-infected mosquitoes into an uninfected population will lead to cytoplasmic incompatibility (Figure 1A). Reduced egg hatch from crosses between infected males and uninfected females favours infected females. For a *Wolbachia* infection to invade an uninfected population, its frequency must exceed a threshold which depends on the fidelity of cytoplasmic incompatibility and maternal transmission and any fitness costs of the infection (O’Neill et al., 1997).

The presence of natural *Wolbachia* infections in a population may result in crossing patterns that make population replacement or suppression more challenging (Figure 1B-E). The following scenarios assume that the natural infection is at fixation in the population. When novel and natural infections are compatible with each other (no reduction in egg hatch in any combination), invasion will depend on the relative fitness of each infection type due to a lack of cytoplasmic incompatibility (Figure 1B). Since transinfections in mosquitoes typically impose fitness costs while natural infections tend to be beneficial (Ross et al., 2019b), population replacement may be unachievable even if high frequencies are reached. In this situation population suppression is impossible due to the lack of cytoplasmic incompatibility in any direction. Such patterns occur in *Cx. pipiens*, with multiple compatible strains coexisting within natural populations (Duron et al., 2011, Atyame et al., 2014).

Incompatibility between males of novel and natural infections and females of the opposite infection type in both directions, or bidirectional cytoplasmic incompatibility, may occur (Figure 1C). Bidirectional incompatibility is desirable for population suppression programs because it reduces the risk that inadvertently released females will replace natural populations (Moretti et al., 2018). Novel *Wolbachia* infections that are bidirectionally incompatible with natural populations have been generated in *Ae. albopictus* (Xi et al., 2006, Calvitti et al., 2010) by first removing the native superinfection which is at high frequency in most natural populations (Kittayapong et al., 2002a, Joanne et al., 2015). Such strains have been deployed successfully for population suppression (Mains et al., 2016). Bidirectional incompatibility can also occur between natural populations of *Drosophila simulans* (O’Neill and Karr, 1990, Montchamp-Moreau et al., 1991), *Nasonia* wasps (Bordenstein and Werren, 2007) and *Cx. pipiens* (Yen and Barr, 1973).

When bidirectional incompatibility occurs, population replacement will be difficult to achieve unless high frequencies of the novel infection are reached. Where population replacement is successful, spread beyond the release area is unlikely since the frequency required for invasion is 50% when fitness is equal (O’Neill et al., 1997). Novel infections may instead persist with natural infections (Telschow et al., 2005), particularly in fragmented populations (Keeling et al., 2003).

Unidirectional incompatibility may also occur between natural and novel infections (Figure 1D-E). If a natural population harbors a double infection and a novel infection with a single *Wolbachia* strain is released, this can result in unidirectional incompatibility favouring the natural infection if one strain of the superinfection is compatible and the other is not (Figure 1D). In this situation, population suppression is impossible and population replacement will be challenging, therefore such infections are not being considered for release. Natural populations of *Ae. albopictus* are superinfected with the *w*AlbA and *w*AlbB strains at a high frequency although either strain may occasionally be lost (Kittayapong et al., 2002a, Joanne et al., 2015), resulting in unidirectional cytoplasmic incompatibility (Dobson et al., 2004).

*Aedes albopictus* with novel *Wolbachia* infections have not been released for population replacement but triple infections may suitable for this purpose (Fu et al., 2010, Zhang et al., 2015). Novel triple infections are unidirectionally incompatible with the natural double infection (Fu et al., 2010, Zheng et al., 2019) (Figure 1E), resulting in a similar pattern to crosses with uninfected mosquitoes (Figure 1A). In cases of unidirectional cytoplasmic incompatibility with the target population (Figure 1A,E), the accidental release of *Wolbachia*-infected females during releases of males for population suppression could lead to population replacement (Dobson et al., 2002). This may be avoided by irradiating release stocks to sterilise any released females, as demonstrated in a recent *Ae. albopictus* population suppression program (Zheng et al., 2019).

Unidirectional cytoplasmic incompatibility can also occur in crosses between two single *Wolbachia* infections (Figure 1D-E) as demonstrated in *Cx. pipiens* (Atyame et al., 2014, Bonneau et al., 2018). In this situation, both strains induce cytoplasmic incompatibility, but one lacks the ability to restore compatibility with males of the other infection. Cytoplasmic incompatibility induction by males is governed by two genes while the ability to restore compatibility by females are governed by a single gene (Shropshire et al., 2018); the two phenotypes are therefore not always linked.

Although natural infections may interfere with releases of novel infections, their presence may also provide opportunities for disease control. *Wolbachia* infections that cause cytoplasmic incompatibility can be released in other locations for population suppression without the need for novel infections (Laven, 1967). Natural infections may also be useful for population replacement if they can block virus transmission (Glaser and Meola, 2010, Mousson et al., 2012).

## Testing a putatively *Wolbachia*-infected laboratory population of *Aedes aegypti*

Of the eight studies reporting natural *Wolbachia* infections in *Ae. aegypti*, only two established laboratory populations (Table 1). We obtained one of these populations with the intention of examining crossing patterns between natural infections and novel infections that are being deployed into the field (Walker et al., 2011, Ant et al., 2018). An *Ae. aegypti* population from Las Cruces, New Mexico, USA was established in the laboratory in September 2018 (Kulkarni et al., 2019) and kindly provided to us by the authors. We received eggs from the third and fourth generations of this population (denoted LC) which were hatched and maintained in our laboratory according to methods described previously (Ross et al., 2017a).

We performed a single cross to test whether *Ae. aegypti* males with the *w*AlbB strain (Xi et al., 2005, Axford et al., 2016) induced cytoplasmic incompatibility with LC females. LC males do not induce detectable cytoplasmic incompatibility with uninfected (Rockefeller strain) females (Jiannong Xu, personal communication). Zero eggs hatched from a cross between *w*AlbB-infected males and LC females (n = 1027 eggs), indicating that the infection is absent, at a low density or is not closely related to the *w*AlbB infection. Due to the absence of *Wolbachia* in the LC strain as detected through molecular analyses (see below), we did not proceed with further crosses.

We used molecular approaches to try and confirm *Wolbachia* infection in the *Ae. aegypti* LC strain. According to the authors, this population harbors a natural *Wolbachia* infection closely related to the *w*AlbB infection from *Ae. albopictus* (Kulkarni et al., 2019). Real-time PCR/high-resolution melt (RT/HRM) assays were performed as previously described (Lee et al., 2012, Axford et al., 2016) using primers specific to the *w*AlbB *Wolbachia* strain as well as *Aedes* and *Aedes aegypti*-specific primers (Appendix 1). We also used a loop mediated isothermal amplification (LAMP) assay which can detect the *w*AlbB infection with high sensitivity (Jasper et al., 2019). Uninfected *Ae. aegypti* originating from Cairns, Australia and *w*AlbB-infected *Ae. aegypti* (Axford et al., 2016) were included as negative and positive controls respectively in each assay. Through these two approaches we did not detect any *w*AlbB infection in 120 mosquitoes (including larvae and adults from both generations) from the LC population (Appendix 2), demonstrating that the LC laboratory population is not infected with *w*AlbB.

To test whether the LC population harbors any *Wolbachia* infection, we performed additional assays with general *Wolbachia* primers. TaqMan probe assays were performed as described previously (Mee et al., 2015), targeting the 16S rDNA (Appendix 1). We also performed conventional PCR with 16S rDNA and *gatB* primers following methods described by the authors of the original study (Kulkarni et al., 2019). Finally, LAMP assays were performed using our protocol (Jasper et al., 2019) but with primers used to diagnose *Wolbachia* infections by the original study (Kulkarni et al., 2019). From analyses of 72 individuals from both generations with the three molecular assays, zero were infected (Appendix 2).

Negative and positive controls were confirmed in all assays. Through these analyses, we demonstrate conclusively that the LC population does not harbor *Wolbachia*. These results conflict with those from the original study (Kulkarni et al., 2019) and more recent tests by the authors where *Wolbachia* is at a high frequency (28/32, 87.5%) in the fourth laboratory generation (Jiannong Xu, personal communication). Although the reason for this conflicting result is unclear, our study emphasizes the need for independent evaluation of *Wolbachia* infections in *Ae. aegypti*.

## Field survey for natural *Wolbachia* infections in *Aedes aegypti*

Teo et al. (2017) detected *Wolbachia* in *Ae. aegypti* from a site in Kuala Lumpur, Malaysia. To further test *Wolbachia* from Kuala Lumpur, we conducted our own sampling, undertaken as part of a release program with the wAlbB *Wolbachia* infection (Nazni et al., 2019). We sampled 693 *Ae. aegypti* from a site in 2013-2014 before *Wolbachia* releases commenced. We also sampled 382 *Ae. aegypti* from July 2017 to September 2018 from a control site where no *Wolbachia* releases were undertaken. Through conventional PCR and RT/HRM assays (described above) we did not detect *Wolbachia* infection in any individual (Appendix 3), in contrast to Teo et al. (2017). Our results are consistent with a global survey of *Ae. aegypti* where no evidence for natural *Wolbachia* infections was found (Gloria-Soria et al., 2018). Below we discuss the limitations of current studies and describe the evidence needed to confirm the presence of putative natural *Wolbachia* infections.

## Limitations of studies to date

Detections of *Wolbachia* in *Ae. aegypti* are accumulating (Table 1) but the evidence is largely molecular, which is insufficient to diagnose an active *Wolbachia* infection (Chrostek and Gerth, 2019). Coon et al. (2016) were the first to report the detection of *Wolbachia* in natural *Ae. aegypti* populations. In this study, *Wolbachia* were found at a low abundance and frequency in Florida, USA through 16S rRNA sequencing, and then characterized with multilocus sequence typing (MLST). Bennett et al. (2019) and Hegde et al. (2018) also detected *Wolbachia* at a low frequency and abundance through 16S rRNA sequencing but these results could not be validated with PCR amplification. These observations may reflect true infections although there are several potential sources of contamination that can cause false positives (discussed in Chrostek and Gerth (2019)).

Several species of filarial nematodes that infect *Ae. aegypti* harbor obligate *Wolbachia* infections from supergroups C and D (Bouchery et al., 2013). Both Thongsripong et al. (2018) and Carvajal et al. (2019) detected *Wolbachia* in *Ae. aegypti* that aligned to supergroup C. Carvajal et al. (2019) observed substantial diversity in *16S* rDNA and *wsp* sequences, with alignments to supergroups A, B, C, D and J. Given that *Wolbachia* from supergroups C, D and J are not known to occur in Diptera, such diversity is likely explained by contamination from other sources. Species misidentification may also cause false positives if one species harbors *Wolbachia* and the other does not. Both Teo et al. (2017) and Carvajal et al. (2019) used identification keys but did not confirm that samples were *Ae. aegypti* with molecular approaches. Since *Ae. aegypti* and *Ae. albopictus* are sympatric in both locations, detections of *Wolbachia* in *Ae. aegypti* could result from species misidentification. Interspecific matings between infected males and uninfected females might also lead to *Wolbachia* being detected in females given that this has been observed at the intraspecific level (A. Callahan and J. Axford, unpublished data). For molecular confirmation of *Wolbachia* infections, appropriate positive and negative controls are needed. Carvajal et al. (2019) used water as a negative control, but this is inadequate because positive detections may be due to amplification of mosquito nDNA. Mosquitoes or other insects of a known infection status, both *Wolbachia*-infected and uninfected, are needed in each assay for confident diagnosis.

Two studies, Balaji et al. (2019) and Kulkarni et al. (2019), established laboratory colonies of *Ae. aegypti* with natural *Wolbachia* infections, allowing for more robust evidence to be collected on infection status. Kulkarni et al. (2019) demonstrate maternal transmission of the natural *Wolbachia* infection; ten offspring selected randomly from a cross between *Wolbachia*-infected females and uninfected males were infected, while none from the reciprocal cross were infected. However, our inability to detect a *Wolbachia* infection in this laboratory population (as discussed above) suggests that this result may not reflect a true infection.

Balaji et al. (2019) provide several lines of evidence for a natural *Wolbachia* infection in *Ae. aegypti* (Table 1), although there are also limitations to this study. The infected laboratory population exhibited a stable infection frequency of ∼80% across four generations, though reciprocal crosses between infected and uninfected populations are needed to confirm maternal transmission. Treatment of the infected population with tetracycline for four consecutive generations removed the *Wolbachia* infection, although the evidence for this provided in the supplementary information lacks controls. Relative *Wolbachia* densities determined by RT/HRM are broadly consistent with natural infections in *Ae. albopictus* where densities can vary across life stages and between sexes (Tortosa et al., 2010, Calvitti et al., 2015). High *Wolbachia* densities in the ovaries are also consistent with a true infection, since maternal transmission requires infection of the germ line (Veneti et al., 2004) but not somatic tissues, although *Wolbachia* often occupy somatic tissues (Dobson et al., 1999). Electron microscopy images show apparent localization of *Wolbachia* to the ovaries, but images are low resolution and there is no clear distinction between *Wolbachia* and organelles as in other recent studies (Li et al., 2017, Leclercq et al., 2016).

## Evidence required to confirm natural *Wolbachia* infections

From the studies discussed above, we believe the evidence is currently insufficient to indicate that *Ae. aegypti* mosquitoes harbor a natural *Wolbachia* infection. We propose three lines of evidence as a minimum requirement for confirming a *Wolbachia* infection in this species: intracellular localization, maternal transmission and removal of *Wolbachia*. Following molecular detection, laboratory populations can be established from larvae, pupae or adults from *Wolbachia*-positive locations to enable further characterization.

Intracellular localization can be demonstrated by visualizing *Wolbachia* within host tissues such as through fluorescence in situ hybridization (FISH) (Moreira et al., 2009). These observations require appropriate controls including separate probes for *Wolbachia* and host and visualization of tissues with the *Wolbachia* infection removed (see below).

Reciprocal crosses between *Wolbachia*-infected and uninfected mosquitoes can be conducted to demonstrate maternal inheritance. In a true natural infection, only offspring from infected mothers are expected to test positive for *Wolbachia*. Maternal transmission may be imperfect, particularly if the infection has a low density in the ovaries (Narita et al., 2007), so sufficient numbers of offspring need to be sampled. Other patterns of inheritance point against a *Wolbachia* infection or may indicate horizontal transmission.

*Wolbachia* infections can be removed from insects through antibiotic or heat treatment (Li et al., 2014). Novel *Wolbachia* infections can be cleared from *Ae. aegypti* with tetracycline added to larval rearing water or sugar solution fed to adults, through rearing larvae at high temperatures, or a combination of approaches (Ross et al., 2017b, Endersby-Harshman et al., 2019). Following removal, which may require multiple generations of treatment, the lack of infection can be confirmed through molecular approaches or by observing intracellular localization.

Together, these experiments should demonstrate conclusively whether the population harbors a *Wolbachia* infection. Following confirmation, additional experiments would likely be worthwhile, as we discuss below.

## Future directions

The confirmation of natural *Wolbachia* infections in *Ae. aegypti* would open avenues for further research, including applications for disease control programs. Laboratory crosses between natural infections and novel infections are needed to test the potential for natural infections to interfere with releases of novel infections. Surveys for natural infections prior to releases of novel infections may inform release strategies, including the choice of *Wolbachia* strain. Effects of natural infections on host fitness, reproduction and vector competence should be evaluated since they may possess properties useful for reducing virus transmission and/or decreasing population size. Genome sequencing may provide insights into their origin. Finally, natural infections could be transferred to other species through microinjection to study their effects in novel hosts and provide further opportunities for disease control.

Although several studies have now claimed to detect *Wolbachia* in natural *Ae. aegypti* populations, the evidence is not compelling. Studies to date have relied mostly on molecular approaches that may be prone to contamination. These results conflict with a growing body of evidence for a lack of infection in this species which includes a comprehensive global survey (Gloria-Soria et al., 2018), monitoring undertaken before releases of novel infections (Hoffmann et al., 2011) and the data presented here. Our inability to detect *Wolbachia* in a putatively infected laboratory population demonstrates the need for more robust evidence when reporting natural *Wolbachia* infections. Although natural *Wolbachia* infections in *Ae. aegypti* may not exist, releases of novel *Wolbachia* infections are continuing to expand, and new target populations should therefore continue to be monitored prior to releases taking place.

## Data accessibility statement

All data are contained within the manuscript and its appendix.

## Author contributions

PAR conceived the study, performed the live mosquito work and drafted the manuscript, AGC, QY and MJ performed the molecular diagnostics on the laboratory population, AKMA, ANA and WAN conducted the field survey, AAH supervised and coordinated the project and all authors contributed to writing and editing the manuscript.

## Acknowledgements

The authors thank Kelly Richardson for assistance with molecular diagnostics, Jiannong Xu and Aditi Kulkarni for providing the *Aedes aegypti* LC strain and Nancy Endersby-Harshman for coordinating the import of the strain into our quarantine facility. AAH was supported by the National Health and Medical Research Council (1132412, 1118640, www.nhmrc.gov.au), the Australian Research Council (LE150100083, www.arc.gov.au) and the Wellcome Trust (108508, wellcome.ac.uk). The funders had no role in study design, data collection and analysis, decision to publish, or preparation of the manuscript.

---

**Appendix 1.**
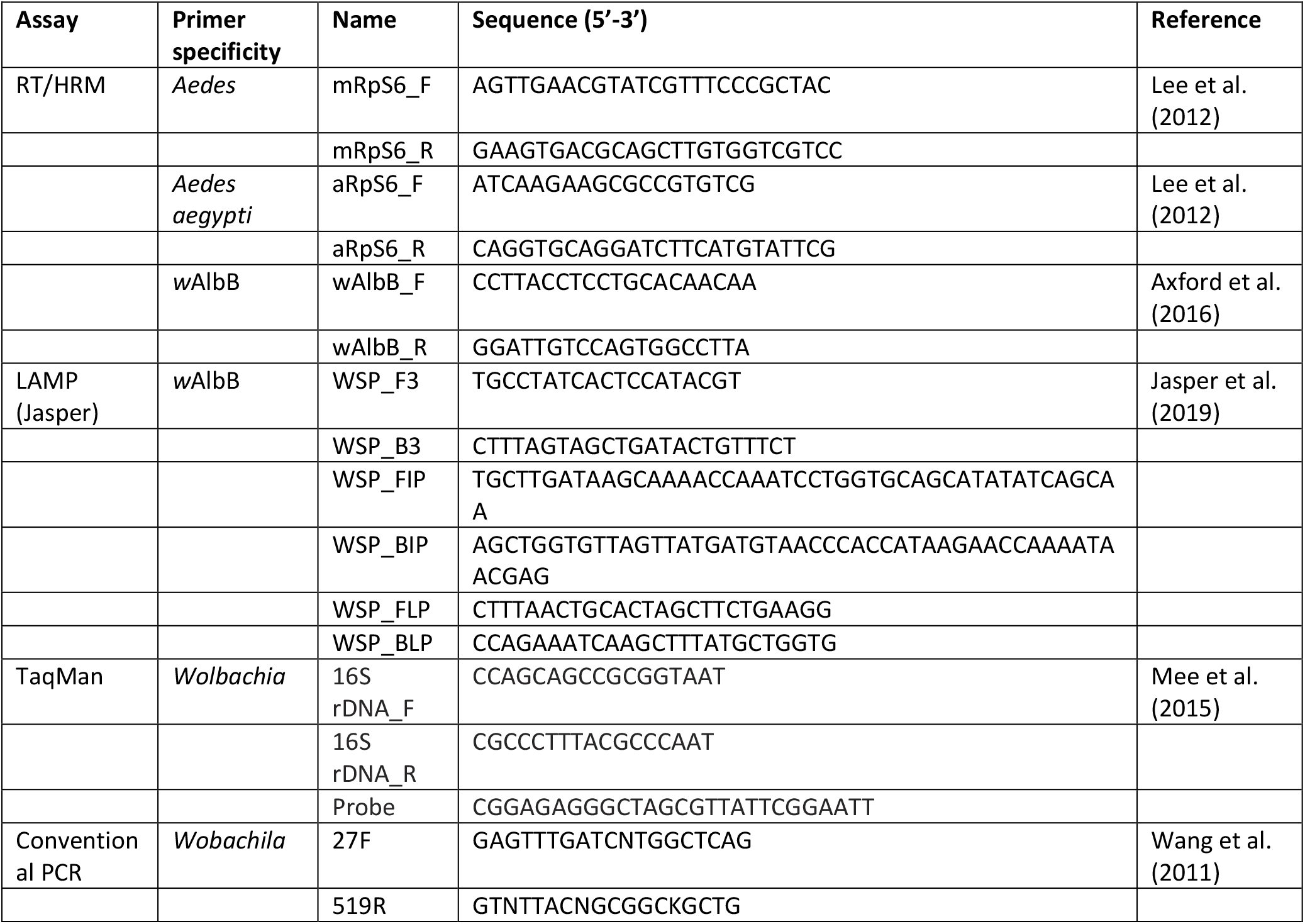
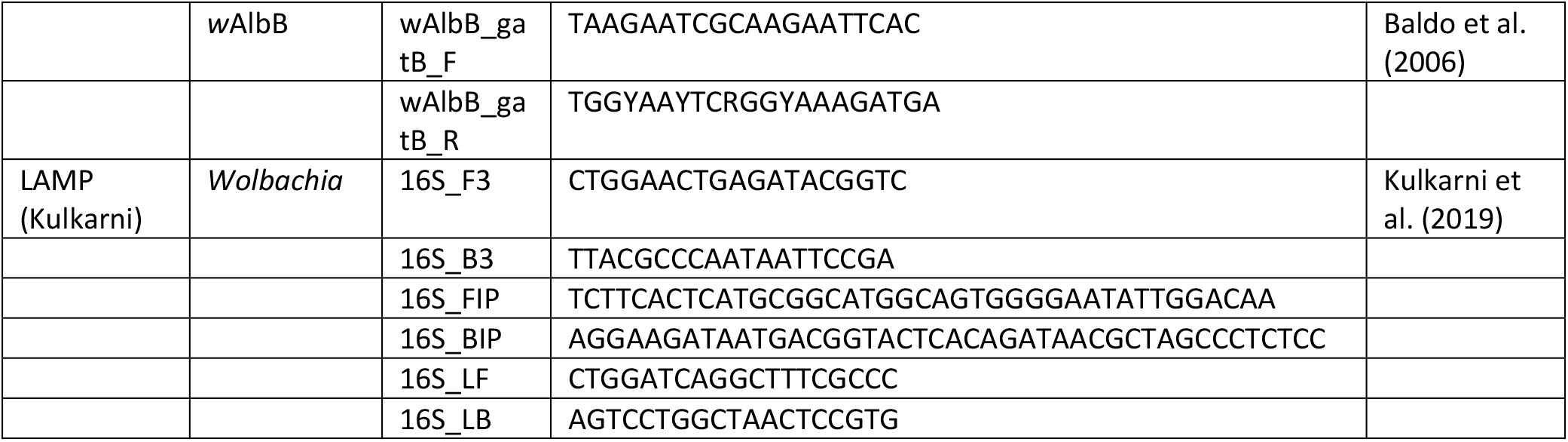
Primers used for detecting *Wolbachia* in the *Aedes aegypti* LC laboratory population with molecular assays.

**Appendix 2.** Molecular detection of *Wolbachia* in the *Aedes aegypti* LC laboratory population.

| Assay | Generation | Life stage | Individuals tested | Pools tested (individuals per pool) | Percent infected |
| --- | --- | --- | --- | --- | --- |
| RT/HRM | 3 | 3<sup>rd</sup> instar larva | 24 | - | 0 |
|  | 4 | 3<sup>rd</sup> instar larva | 24 | - | 0 |
| LAMP (Jasper) | 3 | 3<sup>rd</sup> instar larva | 18 | 3 (6) | 0 |
|  | 3 | Adult | 18 | 3 (6) | 0 |
|  | 4 | 3<sup>rd</sup> instar larva | 18 | 3 (6) | 0 |
|  | 4 | Adult | 18 | 3 (6) | 0 |
| TaqMan | 3 | 3<sup>rd</sup> instar larva | 24 | - | 0 |
|  | 4 | 3<sup>rd</sup> instar larva | 24 | - | 0 |
| Conventional PCR (*Wolbachia*) | 3 | 3<sup>rd</sup> instar larva | 12 | - | 0 |
|  | 4 | Adult | 12 | - | 0 |
| Conventional PCR (wAlbB) | 3 | 3<sup>rd</sup> instar larva | 12 | - | 0 |
|  | 4 | Adult | 12 | - | 0 |
| LAMP (Kulkarni) | 3 | Adult | 12 | 3 (4) | 0 |
|  | 4 | Adult | 12 | 3 (4) | 0 |

**Appendix 3.** Molecular detection of *Wolbachia* in *Aedes aegypti* collected from Ridzuan Condominia, Petaling Jaya (3°04’46.5”N, 101°36’19.9”E) and Section 7, Shah Alam (3°04’10”N, 101°28’58.5”E) in Kuala Lumpur, Malaysia with conventional PCR (Ridzuan Condominia) or RT/HRM (Section 7).

| Location | Date(s) collected | Individuals tested | Percent infected |
| --- | --- | --- | --- |
| Ridzuan Condominia | 2013-2014 | 693 | 0 |
| Section 7 | July 4, 2017 | 22 | 0 |
|  | October 5, 2017 | 5 | 0 |
|  | October 30, 2017 | 27 | 0 |
|  | November 27, 2017 | 5 | 0 |
|  | January 3, 2018 | 5 | 0 |
|  | February 5, 2018 | 26 | 0 |
|  | March 26, 2018 | 45 | 0 |
|  | June 4, 2018 | 8 | 0 |
|  | July 11, 2018 | 44 | 0 |
|  | August 7, 2018 | 118 | 0 |
|  | September 5, 2018 | 77 | 0 |
|  | Total | 1075 | 0 |

